# Aquaporin-4 mis-localization slows glymphatic clearance of α-synuclein and promotes α-synuclein pathology and aggregate propagation

**DOI:** 10.1101/2024.08.14.607971

**Authors:** Molly Braun, Matthew J. Simon, Jay Jang, Keith Sanderson, Justyna Swierz, Mathew Sevao, Alexandra B. Pincus, Allison J. Schaser, Jonathan E. Elliott, Miranda M. Lim, Vivek K. Unni, Abigail G. Schindler, C. Dirk Keene, Caitlin S. Latimer, Jeffrey J. Iliff

**Affiliations:** VISN 20 Northwest Mental Illness Research, Education and Clinical Center (MIRECC), VA Puget Sound Health Care System, Seattle, WA, USA and VA Portland Health Care System, Portland, OR, USA; Department of Psychiatry and Behavioral Sciences, University of Washington School of Medicine, Seattle, WA, USA; Department of Neurosurgery, Medical College of Georgia, Augusta University, Augusta, GA, USA; Department of Anesthesiology and Perioperative Medicine, Oregon Health & Science University, Portland, OR, USA; Neuroscience Graduate Program, Oregon Health & Science University School of Medicine, Portland, OR, USA; Jungers Center for Neuroscience Research, Oregon Health & Science University, Portland, OR, USA; Department of Neurology, Oregon Health & Science University, Portland, OR, USA; Knight Cardiovascular Institute; Oregon Health & Science University, Portland, OR, USA; VISN 20 Geriatric Research, Education and Clinical Center (GRECC), VA Puget Sound Health Care System, Seattle, WA, USA; Department of Medicine, Division of Gerontology and Geriatric Medicine, University of Washington School of Medicine, Seattle, WA, USA; Department of Laboratory Medicine and Pathology, University of Washignton School of Medicine, Seattle, WA, USA; Department of Neurology, University of Washington School of Medicine, Seattle, WA, USA

**Author notes:** **Corresponding Author:** Jeffrey J. Iliff, PhD; **Full address:** VISN 20 MIRECC, VA Puget Sound Healthcare System, 1660 South Columbia Way, Seattle, WA 98108, USA; **E-mail:**.

**Keywords:** Glymphatic, aquaporin-4, AQP4, Astrocytes, Perivascular endfeet, α-syntrophin, α-synuclein, propagation, PFFs, Lewy body dementia

## Abstract

The appearance of misfolded and aggregated proteins is a pathological hallmark of numerous neurodegenerative diseases including Alzheimer’s disease and Parkinson’s disease. Sleep disruption is proposed to contribute to these pathological processes and is a common early feature among neurodegenerative disorders. Synucleinopathies are a subclass of neurodegenerative conditions defined by the presence of α-synuclein aggregates, which may not only enhance cell death, but also contribute to disease progression by seeding the formation of additional aggregates in neighboring cells. The mechanisms driving intercellular transmission of aggregates remains unclear. We propose that disruption of sleep-active glymphatic function, caused by loss of precise perivascular AQP4 localization, inhibits α-synuclein clearance and facilitates α-synuclein propagation and seeding. We examined human post-mortem frontal cortex and found that neocortical α-synuclein pathology was associated with AQP4 mis-localization throughout the gray matter. Using a transgenic mouse model lacking the adapter protein α-syntrophin, we observed that loss of perivascular AQP4 localization impairs the glymphatic clearance of α-synuclein from intersititial to cerebrospinal fluid. Using a mouse model of α-synuclein propogation, using pre-formed fibril injection, we observed that loss of perivascular AQP4 localization increased α-synuclein aggregates. Our results indicate α-synuclein clearance and propagation are mediated by glymphatic function and that AQP4 mis-localization observed in the presence of human synucleinopathy may contribute to the development and propagation of Lewy body pathology in conditions such as Lewy Body Dementia and Parkinson’s disease.

**Summary:** In a human postmortem case series, we observe that neocortical Lewy body pathology is associated with mis-localization of the astroglial water channel aquaporin-4 (AQP4). In mice, α-synuclein is cleared from the brain along perivascular pathways, while loss of perivascular AQP4 localization impairs glymphatic α-synuclein clearance to the CSF. Furthermore, loss of perivascular AQP4 localization promotes the development and propagation of α-synuclein aggregates.

## Introduction

Accumulation of soluble proteins into insoluble aggregates is a defining pathological feature common to a broad range of neurodegenerative diseases including Alzheimer’s disease (AD), Huntington’s disease, amyotrophic lateral sclerosis and Parkinson’s disease (PD). Many of these neurodegenerative diseases feature sleep disruption early in the disease course, and mounting evidence suggests lack of sleep is a biological driver of insoluble protein aggregation^1^. It is thought that intercellular propagation of these aggregates contributes to the stereotyped progressive neuroanatomical spread of pathology that parallels the clinical progression of several neurodegenerative conditions ^2-4^, whereas sleep may serve to enhance clearance of these proteins and prevent aggregation and propagation ^5,6^. This propagation is perhaps most well established among the synucleinopathies, including PD and dementia with Lewy bodies (DLB), and is typified by the age-associated appearance of α-synuclein inclusions in either neuronal or glial cell bodies ^7^. While cell-cell transmission of α-synuclein aggregates has been demonstrated across several model systems ^8-11^, and is modulated by the isoform of aggregated α-synuclein ^12-15^, whether host features within the brain environment, such as AQP4 localization and glymphatic function, alter the the development of α-synuclein pathology or the dynamics of aggregate propagation remains unknown.

Several mechanisms have been proposed to govern intercellular aggregate formation and transmission, including synaptic transmission, extracellular vesicle release and release by damaged or dead cells^16^, however the relative contributions of each to total α-synuclein aggregate propagation is unclear. Common among each of these mechanisms is the obligatory transit of aggregates, either freely floating or enclosed within a vesicle, through the brain interstitium (i.e., extracellular space). The recent characterization of the “glymphatic” pathway, a brain-wide network of perivascular spaces that supports the clearance of interstitial proteins including amyloid β^17^ and tau^18^ through the exchange of cerebrospinal fluid (CSF) and interstitial fluid (ISF) ^17,19,20^, represents one mechanism that may modulate the dynamics of α-synuclein aggregation and extracellular α-synuclein aggregate propagation. Glymphatic function, and the corresponding clearance of these interstitial proteins, increases during sleep, is impaired by acute sleep disruption^5,21^, and is regulated by the circadian cycle^22^. In turn, in human participants, acute sleep disruption impairs the clearance amyloid β and tau from the CSF to the blood^23^, and results in changes in AD biomarkers indicative of pathological burden^24,25^. Sleep disruption more generally is a common early correlate of neurodegenerative conditions^26,27^. Impairment of glymphatic exchange is also observed in aging ^28^, following traumatic brain injury ^18,29^, and in rapid eye movement (REM) sleep behavior disorder (RBD)^27,30^, three additional conditions representing independent risk factors of synucleinopathy^31-38^. Thus, impairment of glymphatic function due to sleep disruption, aging, or following brain injury may promote the process of α-synuclein aggregate propagation.

Perivascular CSF-ISF exchange is is facilitated by the astroglial water channel aquaporin-4 (AQP4), which is localized to perivascular astrocytic endfeet ensheathing the brain microvasculature^17,39^. In mice, deletion of *Aqp4* slows perivascular CSF-ISF exchange and accelerates amyloid β^40,41^ and tau aggregate formation^18,42-44^, while impairment of glymphatic function in aging^28^, TBI^18,29,45^ and ischemic injury^46,47^ are each associated with the loss of perivascular AQP4 localization. In human neuropathological cases, loss of perivascular AQP4 localization is associated with a clinical diagnosis of AD and AD neuropathology^48,49^, while in rodent models loss of perivascular AQP4 localization is sufficient to impair glymphatic function and promote the development of amyloid β pathology^39,41,49^. Yet whether mis-localization occurs in the presence of neocortical Lewy body pathology or whether reduced perivascular AQP4 localization promotes α-synuclein pathological progression have not been directly evaluated.

In the present study we first evaluate whether neocortical Lewy body pathology is associated with perivascular AQP4 mis-localization in a human post mortem case series. We then use a transgenic mouse model to define if loss of perivascular AQP4 localization impairs the perivascular glymphatic clearance of α-synuclein. We last evaluate whether impairment of glymphatic function accelerates the propagation of α-synuclein aggregates in a seeding/propagation mouse model.

## Methods

### Human post-mortem histopathological study

#### Human Post-Mortem Brain Specimens

Brain specimens were acquired from the University of Washington (UW) BioRepository and Integrated Neuropathology (BRaIN) laboratory. Consent for research brain procurement was obtained from the donor or from the legal next of kin, according to the protocols approved by the UW Institutional Review Board. At the time of procurement, the brain was removed in the usual fashion and all brains were extracted, handled, and stored under the same conditions. Briefly, following removal, brains are submersion fixed in formalin for at least two weeks, coronally sectioned at 4mm thickness. Representative regions are then sampled and embedded in paraffin for subsequent histologic assessment. All brains undergo a research diagnostic examination by a board-certified neuropathologist for AD neuropathologic change, Lewy body pathology, and other co-pathologies according to National Institute of Aging-Alzheimer’s Association (NIA-AA) guidelines ^50^. Specifically, for Lewy body disease, immunohistochemical assessments for pathologic α-synuclein in the olfactory bulb, amygdala, brainstem, anterior cingulate cortex, and middle frontal gyrus to identify Lewy bodies and classify the distribution as olfactory bulb only, amygdala predominant, brainstem predominant, limbic, or neocortical^51^. Detailed case information can be found in **Supplemental Table 1**.

#### Case Selection

For human tissue analyses we obtained from the BRaIN lab cases with neocortical Lewy body pathology (n=21, 10 female, 11 male) and sex and age-matched cases without any Lewy body pathology (n=19, 9 female, 10 male). In order to isolate the relationship between neocortical Lewy body pathology and perivascular AQP4 localization from the influence of comorbid amyloid β or tau pathology, we excluded cases with a CERAD score of moderate or severe neuritic plaques and Braak stages V and VI (neocortical involvement).

#### Immunofluorescence for human post-mortem tissue

5 μm thick formalin-fixed, paraffin-embedded tissue samples from frontal cortex were examined from each case. Tissue was deparaffinized, followed by heated antigen retrieval in Tris-EDTA pH 9.0 with tween (Abcam recipe) in a steamer for 20 minutes, followed by further antigen retrieval with 90% formic acid for 4 minutes. Sections were permeabilized in 0.3% Triton X-100 in phosphate buffered saline (PBS) for 15 minutes, rinsed in PBS, and treated with TrueBlack Lipofuscin Autofluorescence Quencher (Biotium, catalog# 23007) to quench lipofuscin and autofluorescence, followed by incubation with rabbit anti-p-S129 (10ug/mL; Abcam; catalog # ab51253) diluted in 5% normal donkey serum in phosphate-buffered saline overnight at 4°C. Following overnight primary incubation, sections were incubated with secondary antibody donkey anti-rabbit Alexa Fluor 647 (1:400; Invitrogen, catalog #A21207) for 2 hours at room temperature. Sections were then incubated with rabbit anti-AQP4 (1:500; Millipore Sigma; catalog #AB3594) and mouse anti-CD31 (1:100; Abcam; catalog #ab9498) diluted in 5% normal donkey serum in PBS overnight at 4°C overnight. Following overnight primary incubation, sections were incubated with secondary antibody donkey anti-rabbit Alexa Fluor 488 (1:500; Invitrogen, catalog # A-21206) and donkey anti-mouse Alexa Fluor 594 (1:200; Invitrogen; catalog #A-21203) for 2 hours at room temperature. Slices were mounted using ProLong™ Diamond Antifade Mountant with DAPI (Invitrogen; catalog #P36971).

#### Quantification of AQP4 and α-synuclein in human tissue

Immunofluorescence imaging was performed on a Keyence BZ-X800 fluorescence microscope with a 20x/0.75 Plan Apo objective. Micrographs were analyzed using FIJI ImageJ software as previously described ^29^, with minor modifications, and the investigators were blinded to case status. For the **laminar shell analysis**, five concentric laminar regions of interest (ROIs) were drawn encompassing the superficial grey matter, middle grey matter, deep grey matter, superficial white matter, and deep white matter. Mean fluorescence intensity was measured for each shell ROI for both AQP4 and p-S129. For area coverage, images were uniformly thresholded to derive the percent of covered area of each ROI. For **line analysis**, linear ROIs were drawn across the tissue, from the pial surface through the grey matter and separate lines were drawn through the white matter to measure changes in AQP4 immunofluorescence intensity. In each section, three linear ROIs were drawn each at the crest of the gyrus, midway down the gyrus, and at the sulcal depth. Replicate ROIs were averaged at each location for each case to reduce signal noise, then averaged across groups. Given the variable distances across the grey and white matter between tissue samples, all gray matter lines were then scaled to a length of 2000 pixels and all white matter lines were scaled to a length of 1000 pixels and fit with a cubic spline function so that they were all the same lengths. For statistical analysis, GM line segments were divided into 3 equal segments corresponding to the superficial, middle, and deep gray matter regions of the laminar shell analysis. WM line segments were divided into 2 equal segments corresponding to superficial and deep white matter. For **analysis of AQP4 localization**, linear radial ROIs were drawn outward from vessel walls, through the surrounding vessel-associated astrocytes and surrounding neuropil. Pixel intensity projections were averaged within regions and within exposure groups to generate the average intensity projections. Vessels were considered to be capillaries if they were less than 10 µm in diameter, while those greater than 10 µm in diameter were considered ‘large vessels.’ Perivascular and non-perivascular segments of these linear ROIs were averaged and plotted. The first 3 pixels were averaged for the perivascular endfoot segment and the last 20 pixels were averaged for the non-perivascular segment. To define the ‘polarization’ of AQP4, we calculated the ratio of perivascular/non-perivascular AQP4 immunofluorescence by dividing the perivascular intensity by the non-perivascular intensities.

#### Statistical Analysis

All human data were analyzed using GraphPad Prism 10 and Python 3.6 software. Two group comparisons were analyzed by two-sided unpaired t-test or Mann Whitney U test, if data was not normally distributed. Multigroup comparisons were made using two-way repeated measures analysis of variance (ANOVA) where appropriate. To examine associations between AQP4 localization and α-synuclein and other variables, we used linear and mixed-effect regressions. Model assumptions were checked and validated. A p value of <0.05 was considered to be significant. Samples sizes were determined by previous experience.

### Rodent studies

#### Materials and antibodies

Soluble α-synuclein (Anaspec, AS-55457) was used at the stock concentration of 1 mg/ml. Mouse wild type sequence preformed-fibrils (PFFs) were generated and prepared as previously described ^11,52^. Primary antibodies included rabbit anti-AQP4 (1:500, Millipore AB3594), rabbit anti-α-syntrophin (1:500, Signalway Antibody 22845), mouse anti-β-actin (1:1000, Novus NB600-501), mouse anti-α-synuclein (1:100, ThermoFisher 32-8100), rabbit anti-phospho-α-synuclein (1:500, Abcam ab51253), mouse anti-GFAP (1:300, Millipore MAB360), mouse anti-NeuN (1:100, Millipore MAB377) and goat anti-Iba1 (1:200, Novus Biologicals NB100-1028). All secondary antibodies for immunofluorescence were generated in donkey, were conjugated to either Alexafluor-488, Alexafluor 594, or Alexafluor 647, used at a concentration of 1:500 and were ordered from ThermoFisher. Hoescht 33342 (1:5000, ThermoFisher H3570) was used to label cell nuclei. CSF quantification of mouse α-synuclein was performed with a commercially available ELISA detection kit (Biomatik EKU07561).

#### Animals

Homozygous *Snta1^-\-^* mice were generated by Dr. Stanley Froehner ^53^ and were obtained from Jackson Laboratories (stock no. 012940). Mice were maintained on a C57Bl6/J background and were homozygously bred. Animals were used between 10-16 weeks of age for all experiments. Age-matched C57Bl6/J mice obtained from Jackson Laboratories were used for wild-type controls. All mice were cared for by the Oregon Health & Science Department of Comparative Medicine in an Association for Assessment and Accreditation of Laboratory Animal Care (AALAC) accredited vivarium. All experiments were performed in accordance with state and federal guidelines and all experimental protocols were approved by the institutional animal care and use committee (IACUC). Power analysis was performed either using pilot experiments or literature review to estimate animal counts for each experiment. Power analyses were run to detect significance levels of p≤0.05, with a power of 80% and effect size of 50%.

#### Immunofluorescence

All tissue slices were generated by 20 µm sectioning on a cryostat. Free-floating immunofluorescence was performed on 20 µm sections for all acute time-frame experiments. PFF experiments immunofluorescence was performed on slide mounted sections. For both free-floating and slide mounted sections, tissue was incubated overnight at 4° C on a rocker with blocking solution (0.3% Triton X-100, 5% normal donkey serum, 2% bovine serum albumin (BSA) in phosphate buffered saline (PBS). Both primary antibody and secondary antibodies were diluted in blocking solution and incubations were performed overnight at 4° C on a rocker. Hoescht staining was applied for 10 minutes during the final set of washes to label nuclei. Floating sections were then positioned directly onto coverslips with a paintbrush. All samples were mounted with Fluoromount-G mounting media (Southern Biotech 0100-01) or homemade Mowiol 4-88.

#### Protein isolation and Western blot

Whole brain tissue lysates were generated by dounce homogenization in RIPA buffer (ThermoFisher 89900) with added protease inhibitor cocktail (Sigma-Aldrich 11836170001). Samples were spun at 2000g for 10 minutes at 4° C to remove cellular debris then frozen at - 80° C. Protein concentration was quantified by Pierce BCA protein assays (ThermoFisher 23225). Western blots were run on NuPage 4-12% Bis-tris gels. 50µg of protein were combined with LDS sample buffer and run on the gel for 90 minutes at 200mV on ice at 4°C. Protein transfer was done using Immobilon-FL membranes and was run at 30mV for 60 minutes at 4°C. Primary and secondary antibodies were applied over the course of 3 hours using the Invitrogen iBind Flex system. Bands were imaged using a Licor Odyssey CLx fluorescence gel imaging system. Values for each sample were generated by averaging across two independent blots. Abundance was determined by measuring the abundance relative to the loading control band (β-ACTIN).

#### Anesthesia

For all acute time-course experiments, anesthesia was briefly induced under isoflurane (2-4%), then maintained under ketamine/xylazine (KX) anesthesia for the duration of the experiment (0.10 mg/g ketamine, 0.01 mg/g xylazine, i.p.). For chronic experiments (e.g., PFF-injections), mice were induced and maintained under isoflurane anesthesia. During all surgeries, mouse body temperature was maintained by heating pads and overhead heating lamps. For long surgeries, (90-120 minutes) mice were given intraperitoneal saline approximately halfway through the surgical procedure to minimize dehydration.

#### Intraparenchymal injections

Intracortical injections targeted the motor cortex (1.0 mm, -1.5 mm, 0.3mm, relative to bregma). Intrastriatal injections targeted caudate putamen (2.8 mm, -0.5 mm, 3.5 mm, relative to bregma). Anesthetized mice were head-fixed in a stereotaxic frame. Fur was cleared from the skin above the skull using Nair. An incision was made along midline to expose the skull and connective tissue was removed with a cotton-tipped applicator. A stereotax-mounted dremel was used to create a small burr-hole at the medial/lateral and anterior/posterior bregma coordinates. The dremel was replaced with a 10µl Hamilton syringe outfitted with a 26G needle (Hamilton 80334) that was loaded into a syringe pump and backfilled with either a 10kD dextran + α-synuclein cocktail or PFFs. The needle tip was then lowered slowly into the brain through the burr hole to the depth of the dorsal/ventral bregma coordinates. The needle was left in place for 1 minute prior to the start of infusion. PFFs were infused at a rate of 200 nL/minute for 12.5 minutes for a final volume of 2.5µl. The needle was then left in the brain for 3 minutes to prevent reflux of PFFs up the needle tract, then extracted and the skin was sutured over the injection site. For the α-synuclein injections, infusions were performed at a rate of 50 nL/minute for 20 minutes for a final volume of 1µl to minimize pressure-driven bulk flow of tracer. The needle was left in place and anesthesia depth was monitored and adjusted as necessary (as described above) over the course of the experimental time frame (120 minutes). Upon completion of both experiments, transcardiac perfusion with 4% paraformaldehyde (PFA) was performed under anesthesia. For the intrastriatal injections, a solution of the lipophilic dye DiD was perfused prior to PFA to label large blood vessels as previously described ^54^. For the PFF dataset, CSF was extracted immediately prior to perfusion (see below). Brains were post-fixed overnight in PFA, followed by an overnight incubation in 30% sucrose to cryo-protect the tissue. Brains were then frozen in OCT and stored at -80°C until sectioning.

#### CSF collection and ELISA assays

Both the basal and post-PFF CSF α-synuclein levels were quantified using a commercially available enzyme-linked immunosorbent assay (ELISA). For CSF collection, a 30 gauge needle was attached to PE-10 tubing and connecting it to a Hamilton syringe in a syringe pump (no backfill). The atlanto-occipital membrane was exposed in isoflurane (1-3%) anesthetized mice and the needle was inserted into the cisterna magna. The syringe pump was then run-in reverse at a speed of 0.5 µL/minute until the AOM had completely collapsed around the needle tip (after approximately 7-8 µL had been collected. Animals were excluded if obvious leakage of CSF occurred during needle placement, or if blood was detected in the sample (as observed by eye before and/or after centrifugation of the CSF). Based on these criteria, four mice were excluded in the endogenous CSF expression study, and one mouse was excluded in the PFF CSF analysis. ELISA assay was performed according to manufacturer instructions. Due to limited sample quantity and low anticipated protein abundance, two technical replicates were run per sample.

#### Confocal and widefield fluorescence imaging

Full-brain imaging was performed using an automated, slide-scanning confocal microscope (Zeiss AxioScan.Z1). Images were acquired using a 20X 0.8 PlanApo objective. High-magnification images were acquired using an inverted laser-scanning confocal microscope equipped with an AiryScan super-resolution detector (Zeiss LSM 880 with Fast Airyscan). Images were collected using either a 63X 1.4 Plan Apo objective or a 20X Plan Apo objective. Based on the semi-quantitative nature of the analysis, all images were acquired to minimize signal outside of the dynamic range of the detector.

#### Image and Statistical Analysis

All image analysis was performed using Image J FIJI software ^55^. All statistical analysis was performed using Graphpad Prism 7. Linear adjustments to minimum and maximum histogram values were made to many of the images presented in the manuscript to improve clarity, however all image analysis was performed on images as acquired, or uniformly size-scaled images (due to computer memory limitations). Due to the automated acquisition of large-scale images, a small percentage (1-2 images across all datasets) exhibited poor focusing and were thus excluded from further analysis. Analyses requiring subjective distinctions (ROI generation, cell counts, threshold identification) were performed in a blinded, manual fashion. During this process, ventricles as well as obvious imaging artifacts were excluded from regional and total brain ROIs. FIJI scripts were generated to automate all image measurements once ROIs and/or thresholds were established. Area coverage was calculated using a threshold-based approach. Briefly, a uniform threshold was established prior to analysis that most accurately captured “real” fluorescent signal for any given tracer or immunolabel. A percentage of the total ROI covered by the threshold generated ROI was then calculated. For the PFF dataset, due to intermittent tissue drying artifacts, thresholds were adjusted to best reflect P-syn-specific signal in a blinded fashion. Furthermore, to most accurately capture the punctate nature of the aggregates, an additional analysis using the “analyze particles” function in FIJI was used to generate an aggregate count. All plots in the figures are presented with bars representing mean ± S.E.M. Mann-Whitney U tests were used to compare all datasets with only 2 values as many exhibited non-parametric distributions. All statistical tests are also listed in the text and figure legends.

## Results

### Neocortical α-synuclein pathology in the human post-mortem brain is associated with loss of perivascular AQP4 polarization

We assessed AQP4 expression and localization in frontal cortex from a case series of human post-mortem brain tissue from the University of Washington Precision Neuropathology Core. Participants either had neocortical Lewy bodies (‘Case’, n=21, 10 female, 11 male) or no Lewy body pathology (‘Control’, n=19, 9 female, 10 male), as determined by standard neuropathological evaluation. Summarized demographic information for all participants is provided in **Table 1**, while individual case information is provided in **Supplemental Table 1.** Overall, participant ages ranged from 62 to 100 years old, with a mean age of 83.75 and median age of 85. Cases and Controls were well matched in terms of age, sex and comorbid tau pathology (assessed by Braak stage). We performed immunofluorescent triple labeling of AQP4, phosphorylated α-synuclein (s129, P-syn), and an endothelial marker CD-31. Representative fluorescence micrographs of frontal cortical gyrus, glia limitans, and vessels from control and Lewy body positive cases are shown in **Figure 1a-d**. We observed increased AQP4 immunofluorescence at the pial surface of Lewy body-positive cases compared to the Lewy body-negative controls (**Figure 1a-b**). This was illustrated with quantification of linear AQP4 immunofluorescence intensity plots across the tissue. Linear ROIs spanning the pial surface through the cortical gray matter (GM) and white matter (WM) were used to examine macroscopic changes in AQP4 immunofluorescence. These ROIs were drawn at the crest of the gyrus (shown in **Figure 1e**), midway down the gyrus, and at the sulcal depth. The immunofluorescence intensity for each participant was averaged and compared across groups. For AQP4 immunofluorescence, when the linear ROIs were divided further into segments corresponding to the laminar shell regions, we detected main effects of depth (p<0.0001) and depth x group interactions (p=0.0197) in the superficial GM; main effects of depth (p<0.0001) in the middle GM and superficial WM; and a main effect of depth x group interaction (p<0.0001) in the deep GM. Complete fluorescence intensity linear ROIs are shown in **Supplemental Figure 1**.

**Figure 1:**
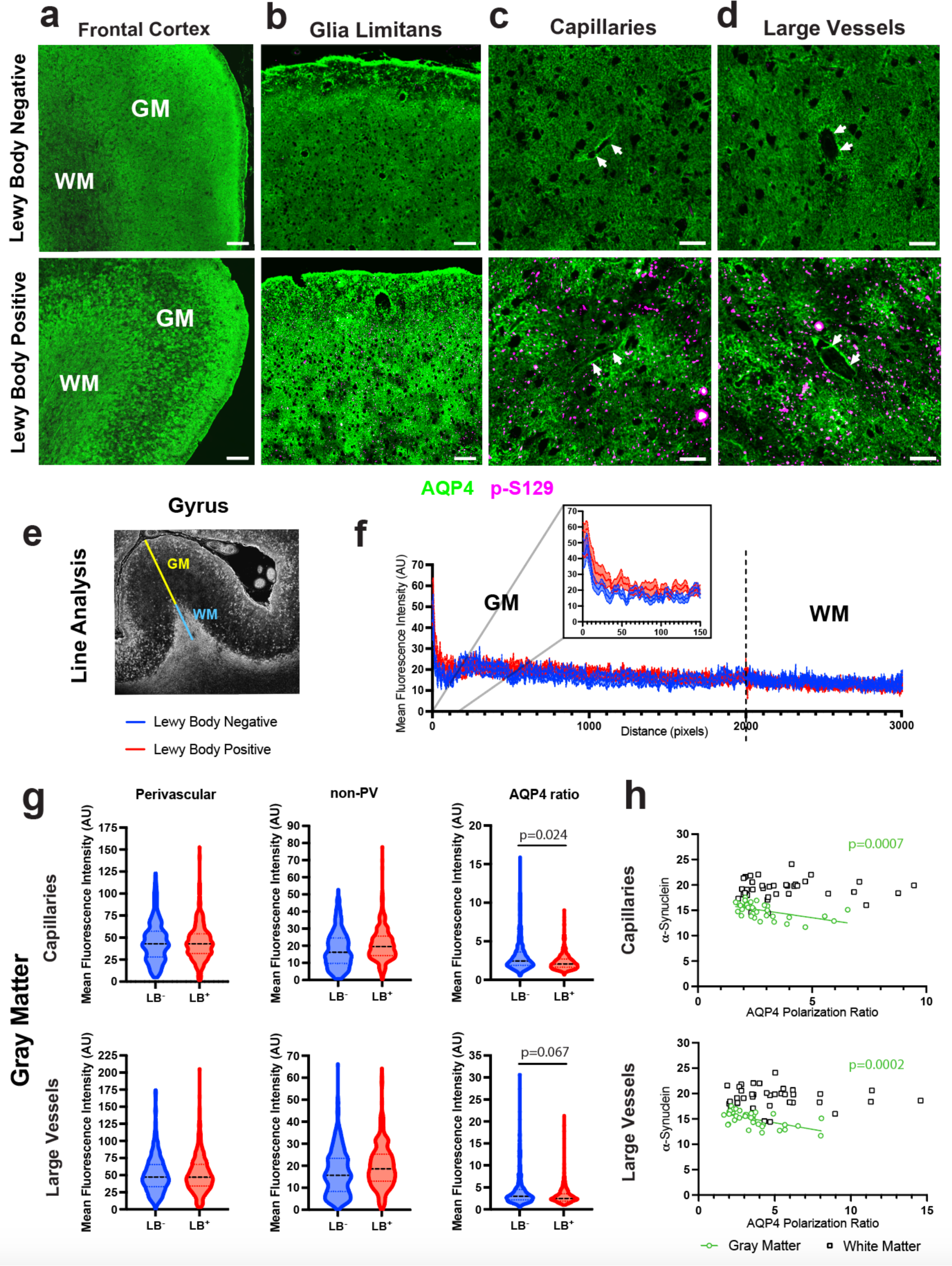
α-Synuclein pathology is associated with decreased AQP4 polarization in the gray matter of human post-mortem brain tissue. (**a**) Representative micrographs of AQP4 and phosphorylated α-synuclein (p-S129) immunoreactivity in the frontal cortex of human post-mortem brain with and without Lewy Body disease. Scale bars = 500 µm (**b)** Higher resolution representative micrographs of the glia limitans. Scale bars = 100 µm. (**c**) capillaries, and (**d**) large vessels. Scale bars = 50 µm (**e**) Representative image of drawn line ROIs extending from the cortical surface through the gray matter (GM) and through the subcortical white matter (WM) that were used to generate intensity projection plots of AQP4 across the tissue. (**f**) Averaged intensity plots of immunofluorescence intensity at the peak of the gyrus are shown for 3 GM/3 WM lines per subject for n=19-21 subjects per group. We detected main effects of depth (p<0.0001) and depth x group interactions (p=0.0197) in the superficial GM, main effects of depth (p<0.0001) in the middle GM and superficial WM, and a main effect of depth x group interaction (p<0.0001) in the deep GM for AQP4 immunofluorescence. **(g)** Quantification of binned AQP4 intensity plot segments surrounding capillaries and large vessels in the GM. Linear ROIs were drawn through the perivascular endfoot, continuing through the surrounding astrocytes and parenchyma to measure AQP4 localization. Both intensity projection plots were averaged for a single intensity plot per vessel. For graphing of perivascular and non-perivascular segments, the first 3 pixels were averaged for the perivascular endfoot segment and the last 20 pixels were averaged for the non-perivascular segment. In the violin plots, the central line represents the median, upper line represents the upper interquartile range (IQR), and the lower line represents the lower IQR. AQP4 was measured surrounding an average of 10 vessels per type per region per subject, for a total of n=523-534 capillaries and n=535-540 large vessels assessed throughout the GM from n=19-21 subjects per group. Random effects mixed effects analysis was performed, to control for sex, age, and correlation within subject (refer to Table 2 for further details), finding a significant decrease in AQP4 polarization ratio surrounding gray matter capillaries (p=0.024) and a trend in decreased AQP4 polarization ratio surrounding gray matter large vessels (p=0.067). (**h**) Linear regression analyses of AQP4 localization and p-S129 showed a negative relationship between AQP4 polarization ratio and α-synuclein pathology surrounding capillaries (p=0.0007) and large vessels (p=0.0002) in the GM from n=19-21 subjects per group.

**Table 1:** Summary of demographic information for participants in post-mortem histopathological study.

| **Frontal Cortical Samples** |  |  |  |  |  |  |
| --- | --- | --- | --- | --- | --- | --- |
|  | **n** | **Age** | **Sex (M/F)** | **Lewy Body Diagnosis** | **Braak Stage** | **CERAD** |
| **Control** | 19 | 86<br>[77, 100] | 10/9 | None | II<br>[I, IV] | Sparse (1)<br>[0, 1] |
| **Case** | 21 | 84<br>[62, 98] | 11/10 | Neocortical | III<br>[I, IV] | Sparse (1)<br>[0.5, 1] |

Values are expressed as median [interquartile ranges].

**Table 2:** Linear mixed effects models, with random effect for participant, for perivascular AQP4, non-perivascular AQP4, and AQP4 polarization ratio surrounding gray matter capillaries and large vessels.

| Capillaries: Perivascular AQP4 |  |  |  |  |
| --- | --- | --- | --- | --- |
| Predictors | Coefficient | 95% Confidence Interval | z | p |
| Intercept | 19.786 | [-31.005, 70.577] | 0.7464 | 0.445 |
| Sex | 3.051 | [-7.088, 13.190] | 0.590 | 0.555 |
| Age | 0.289 | [-0.330, 0.908] | 0.915 | 0.360 |
| Lewy Body Status | -1.688 | [-10.475, 7.099] | -0.376 | 0.707 |
| Capillaries: Non-Perivascular AQP4 |  |  |  |  |
| Predictors | Coefficient | 95% Confidence Interval | z | p |
| Intercept | -2.703 | [-29.060, 23.654] | -0.201 | 0.841 |
| Sex | 3.046 | [-2.224, 8.315] | 1.133 | 0.257 |
| Age | 0.231 | [-0.090, 0.553] | 1.413 | 0.158 |
| Lewy Body Status | 1.528 | [-3.039, 6.095] | 1.413 | 0.512 |
| Capillaries: AQP4 Polarization Ratio |  |  |  |  |
| Predictors | Coefficient | 95% Confidence Interval | z | p |
| Intercept | 6.513 | [3.126, 12.031] | 3.395 | **0.001** |
| Sex | -0.385 | [-1.276, 0.504] | -1.008 | 0.313 |
| Age | -0.037 | [-0.098, 0.010] | -1.588 | 0.112 |
| Lewy Body Status | -0.746 | [-1.616, -0.074] | -2.256 | **0.024** |
| Large Vessels: Perivascular AQP4 |  |  |  |  |
| Predictors | Coefficient | 95% Confidence Interval | z | p |
| Intercept | 25.614 | [-28.129, 79.357] | 0.934 | 0.350 |
| Sex | 3.188 | [-7.567, 13.944] | 0.581 | 0.561 |
| Age | 0.291 | [-0.363, 0.946] | 0.872 | 0.383 |
| Lewy Body Status | -0.826 | [-10.137, 8.486] | -0.174 | 0.862 |
| Large Vessels: Non- Perivascular AQP4 |  |  |  |  |
| Predictors | Coefficient | 95% Confidence Interval | z | p |
| Intercept | 4.180 | [-22.179, 30.540] | 0.311 | 0.756 |
| Sex | 4.024 | [-1.257, 9.304] | 1.493 | 0.135 |
| Age | 0.134 | [-0.187, 0.455] | 0.818 | 0.413 |
| Lewy Body Status | 1.739 | [-2.836, 6.313] | 0.745 | 0.456 |
| Large Vessels: AQP4 Polarization Ratio |  |  |  |  |
| Predictors | Coefficient | 95% Confidence Interval | z | p |
| Intercept | 8.688 | [2.929, 14.448] | 2.957 | **0.003** |
| Sex | -0.323 | [-1.474, 0.828] | -0.550 | 0.582 |
| Age | -0.052 | [-0.122, 0.018] | -1.459 | 0.145 |
| Lewy Body Status | -0.932 | [-1.928, 0.063] | -1.835 | 0.067 |

Next, we evaluated AQP4 immunofluorescence intensity and localization at the level of perivascular astroglial endfeet surrounding capillaries (**Figure 1c**) and large vessels (**Figure 1d**) throughout the gray matter (**Figure 1g**). To quantify perivascular AQP4 expression and localization, linear radial ROIs were drawn outward from the vessel walls, through the surrounding astrocytes and neuropil (**Supplementary Figure 2a, 3a**). Linear ROIs were averaged within regions to generate the average intensity projections shown. Vessels were considered to be capillaries if they were less than 10 µm in diameter, while those greater than 10 µm in diameter were considered ‘large’ vessels. Perivascular and non-perivascular segments of these linear ROIs were averaged and plotted. To define the perivascular ‘polarization’ of AQP4, we calculated the ratio of perivascular/non-perivascular AQP4 immunofluorescence. We used linear random effects mixed models (with random effect for participant), controlling for age and sex, to define the relationship between dichotomous Lewy body status and perivascular AQP4 polarization across all vessels evaluated. Full model outputs are provided in **Table 2**. The distributions of capillary and large vessel perivascular and non-perivascular AQP4 immunofluorescence intensity, and the perivascular AQP4 polarization ratio are provided in **Figure 1g**. The P values for the Lewy body status term of the linear random effects mixed model is provided in the plots for the perivascular AQP4 polarization ratios. In the GM, perivascular AQP4 fluorescence surrounding capillaries and large vessels did not differ between Lewy body-positive cases and controls. Non-perivascular AQP4 immunofluorescence tended to be higher in Lewy body-positive cases, resulting in a significant decrease in perivascular AQP4 polarization ratio surrounding capillaries (p=0.024) and a trend towards reduced perivascular AQP4 polarization surrounding large vessels (p=0.067) throughout the GM (**Figure 1h**). Full model outputs for linear random effects mixed models (with random effect for participant), controlling for age and sex, for gray matter capillaries or large vessels can be seen in **Table 2**. In general, there was no significant effect of age or sex on AQP4 expression or perivascular AQP4 localization. More laminar characterization of AQP4 localization in the Superficial GM, Middle GM, Deep GM, Superficial WM, and Deep WM surrounding capillaries and large vessels can be seen in **Supplementary Figures 2 and 3**, respectively. In general, increased non-perivascular AQP4 localization, resulting in decreased AQP4 polarization ratio, was observed throughout the GM but not the WM. We also evaluated immunofluorescence and area coverage of P-syn and AQP4 and across five concentric laminar ROIs above (**Supplementary Figure 4a**). As expected, P-syn was significantly increased in the Middle GM (p=0.0089), Deep GM (p=0.0039), superficial WM (p=0.0087), and deep WM (p=0.0140) in Lewy body-positive cases compared to Lewy body-negative controls, while macroscopic AQP4 immunofluorescence did not differ between groups (**Supplementary Figure 4b-f)**.

To more granularly define the relationship between neocortical P-syn levels and perivascular AQP4 localization, we conducted linear regression analysis. We observed that decreasing perivascular AQP4 ratio was significantly associated with increasing P-syn immunofluorescence throughout the GM for both capillaries (p=0.0007, R^2^=0.2886) and large vessels (p=0.0002, R^2^=0.3322) (**Figure 1h**). Using this approach in each cortical GM laminae, we observed that decreasing perivascular AQP4 ratio was significantly associated with increasing P-syn immunofluorescence throughout the Superficial GM (p=0.0212, R^2^=0.1465; p=0.0150, R^2^=0.1618), Middle GM (p=0.0266, R^2^=0.1403; p<0.0390, R^2^=0.1303), and Deep GM (p=0.0023, R^2^=0.2419; p=0.0071, R^2^=0.1943) for both capillaries and large vessels, respectively (**Supplementary Figure 5**). No such association was observed in the Superficial WM and Deep WM layers. In total, these data demonstrate that reduced gray matter perivascular AQP4 polarization ratio is associated with neocortical Lewy body pathology in human post mortem frontal cortical tissue.

### Perivascular glymphatic clearance of a-synuclein is impaired in the Snta1^-/-^ mouse

In two pior human neuropathological case series, we have reported that frontal cortical perivascular AQP4 polarization is reduced with aging, and in the presence of Alzheimer’s disease pathology ^48,49^. AQP4 maintains its perivascular localization through its association with the dystrophin-associated complex, which includes the adaptor protein α-syntrophin (SNTA1) ^53,56^. Genetic deletion of *Snta1* in mice abolishes the perivascular localization of AQP4, although expression of AQP4 remains largely unchanged ^49^ (**Figure 2a-d**). Glymphatic function is impaired in *Snta1^-/-^* compared to wild type mice ^39,49^, and deletion of *Snta1* exacerbates amyloid β pathology in two different mouse models of amyloidosis ^41,49^.

**Figure 2:**
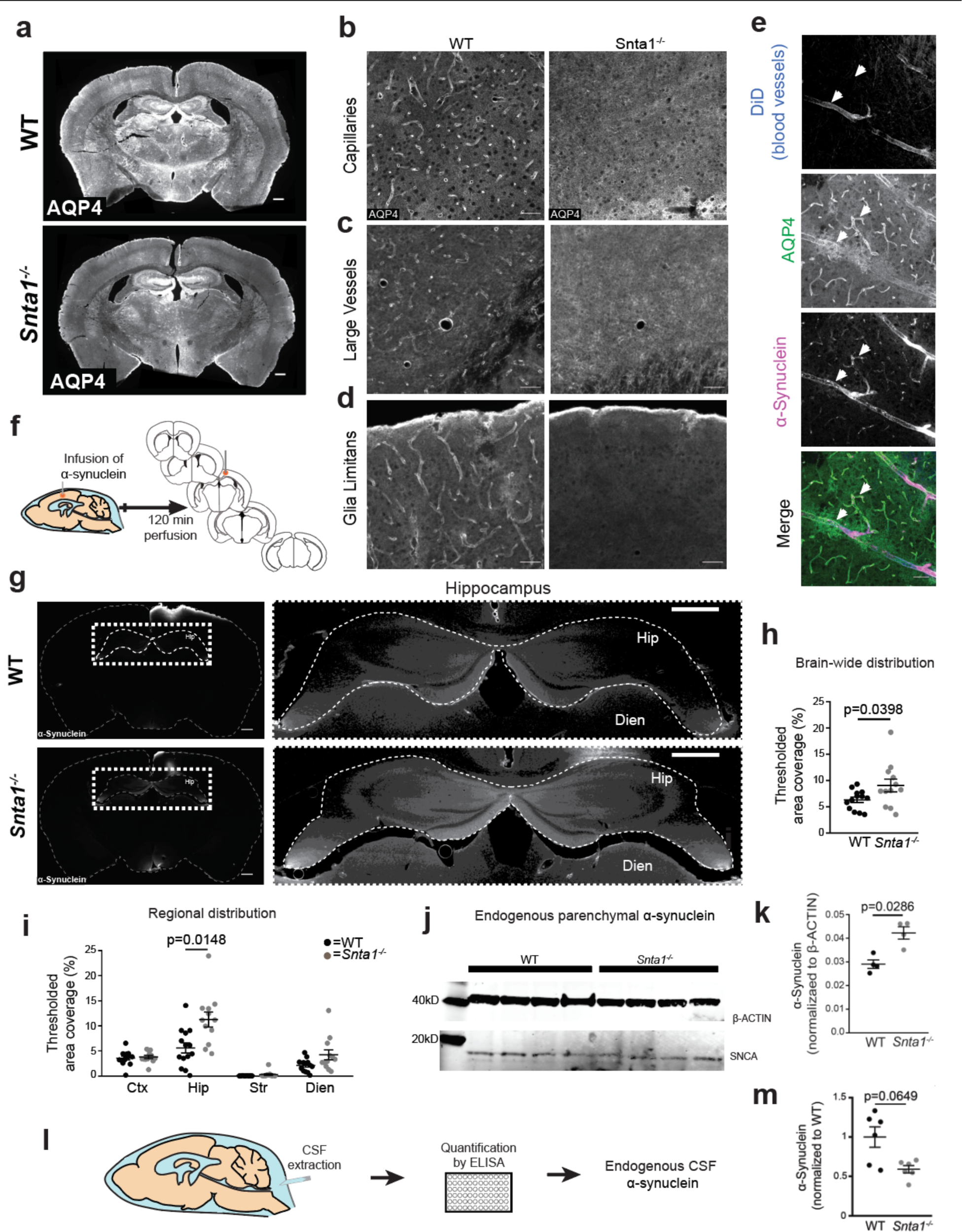
Deletion of α-syntrophin eliminates perivascular localization of AQP4 and slows interstitial α-synuclein clearance. (**a**) Immunofluorescence labeling of AQP4 reveals the robust loss of perivascular localization in *Snta1^-/-^* mice. Scale bars = 500 µm. (**b**) Representative AQP4 labeling surrounding capillaries (**c**) large vessels, and at the (**d**) glia limitans. (e) 30 minutes following intraparenchymal injection of soluble α-synuclein (magenta) in wild type mice, preferential accumulation is observed along the perivascular space of both capillaries and arterioles (arrows). Vessels labeled by vascular infusion of the lipophilic dye DiD (blue), outer boundary of perivascular space identified by immunofluorescence labeling of Aqp4 (green). (**f**) Schematic representation of α-synuclein efflux studies following motor cortex infusion. The five sections in which distribution was evaluated are shown at the right. (**g**) Representative images demonstrating brain-wide α-synuclein distribution 120 minutes post injection Scale bar = 500 µm. Insets at right contain higher magnification images of hippocampus (outlined) with linearly scaled fluorescence. Scale bar = 500 µm. (**h**) Total area coverage by α-synuclein integrated across all 5 sections indicates elevated levels of α-synuclein in *Snta1^-/-^*mice 120 minutes after injection (p = 0.0398, Mann-Whitney U test, n=12-15/group) (**i**) Sub-regional α-synuclein area coverage assessment for cortex (CTX), hippocampus (HIP), striatum (STR) and diencephalon (DIEN). Statistically significant differences are observed in hippocampus (p_adj_=0.0148, multiple t-tests with Holm-Sidak correction, n=12-15/group) (**j**) Representative western blot used to quantify SNCA and β-ACTIN. (**k**) Quantification of western blot, showing increased endogenous parenchymal α-synuclein in *Snta1^-/-^* mice (p=0.0286, Mann-Whitney U test, n=4/group). (**l**) Schematic representation of collection of CSF and ELISA-based quantification. (**m**) Endogenous CSF α-synuclein concentrations following CSF extraction reveals a trend towards reduced CSF α-synuclein in *Snta1^-/-^* mice (p=0.0649, Mann-Whitney U test, n=5-6/group). Raw α-synuclein concentrations are normalized to the mean wild-type CSF α-synuclein concentration. For all plots, black bars represent the mean ± S.E.M.

We next tested whether loss of perivascular AQP4 localization in the *Snta1^-/-^* mouse impaired the perivascular glymphatic clearance of α-synuclein from the brain insterstitium to the CSF. We first evaluated whether α-synuclein is cleared from the brain along perivascular pathways. Animals were injected intra-parenchymally with fluorescently-tagged α-synuclein and brains were fixed 30 minutes later. Immunofluorescence demonstrated that fluorescently-tagged α-synuclein preferentially accumulates along the cortical microvasculature, labeled with DiD (injected iv prior to perfusion to label blood vessels) and AQP4 (to label perivascular astroglial endfeet) (**Figure 2e**). To evaluate whether α-synuclein efflux is dependent upon perivascular AQP4 localization, fluorescently-tagged α-synuclein distribution was evaluated 120 min after intracortical injection in wild type and *Snta1*^-/-^ mice (**Figure 2f-i**). Parenchymal α-synuclein fluorescence intensity levels, when integrated across all regions, were higher in the *Snta1*^-/-^ compared to wild type mice (**Figure 2g, 2h** p=0.040; n=12-13/group), reflecting greater retention of the injected fluorescent α-synuclein. This change was driven primarily by significantly higher levels of α-synuclein in the hippocampus (**Figure 2i** p_adj_=0.015) and levels that tended to be higher in the diencephalon (p_adj_ = 0.106) of *Snta1*^-/-^ compared to wild type mice.

To assess if loss of perivascular AQP4 localization reduces the glymphatic clearance of endogenous α-synuclein, we measured soluble α-synuclein levels in the brain parenchyma and CSF of *Snta1*^-/-^ compared to wild type mice. Endogenous α-synuclein levels were measured in the brain tissue of wild type and *Snta1*^-/-^ mice by Western blot, and in the CSF by ELISA. Consistent with retention of interstitial solutes including α-synuclein (**Figures 2f-i**), parenchymal soluble α-synuclein levels were significantly increased (**Figure 2j, 2k**, p=0.0286, n=4/group) in *Snta1^-/-^*mice compared to wild type mice, while CSF α-synuclein levels were 41% lower in *Snta1^-/-^* mice compared to wild type mice (**Figure 2l, 2m,** p=0.065, n=5-6/group).

### Exacerbated α-synuclein aggregation in Snta1^-/-^ mice

To evaluate whether loss of perivascular AQP4 localization and impaired perivascular CSF-ISF exchange exacerbates pathogenic α-synuclein aggregation and propagation, we utilized a mouse model of synucleinopathy propagation involving a single intracortical inoculation with synthetic α-synuclein pre-formed fibrils (PFFs, **Figure 3a**) ^11^. Consistent with the initial description of this model, four months following injection of PFFs into the motor cortex of wild type mice, widespread P-syn immunoreactivity was detectable in ipsilateral cortex and hippocampus as well as more distal areas including the contralateral cortex, hippocampus and striatum (**Figure 3b**). Immunofluorescence revealed that as previously reported ^11^, P-syn co-localized with the neuronal nuclei marker NeuN (**Supplemental Figure 6a**). Both diffuse P-syn immunoreactivity and dense inclusion bodies were detected in neurons. Unexpectedly, P-syn also co-localized with markers of astrocytes (GFAP) and microglia (Iba1), which do not express high levels of endogenous α-synuclein ^57,58^ (**Supplemental Figure 6a-b**). In *Snta1*^-/-^ mice injected with PFFs, though the general anatomical distribution of P-syn immunoreactivity did not differ from that in wild type animals, the numbers of aggregates were markedly increased in *Snta1*^-/-^ mice (**Figure 3b-c**, p=0.008, n=5/group). This increase was greatest in the cortex (p_adj_=0.109), but was also observed across the striatum, hippocampus and diencephalon (p_adj_=0.200). CSF α-synuclein was also evaluated by ELISA in these animals immediately prior to sacrifice. In agreement with the baseline CSF α-synuclein differences reported above (**Figure 2k**), we observed that CSF levels of soluble α-synuclein were 32% lower in *Snta1*^-/-^ mice than in wild type animals, 4 months after PFF injection (**Figure 3d**, p=0.064, n=4-5/group). When CSF α-synuclein levels were plotted against parenchymal P-syn immunoreactivity for the wild type (n=5) and *Snta1^-/-^* mice (n=4; due to failed CSF sampling in one animal), a strong significant negative association was observed (**Figure 3e**, p=0.005, R^2^=0.706), with lower CSF α-synuclein levels corresponding to greater P-syn immunoreactivity across all animals.

**Figure 3:**
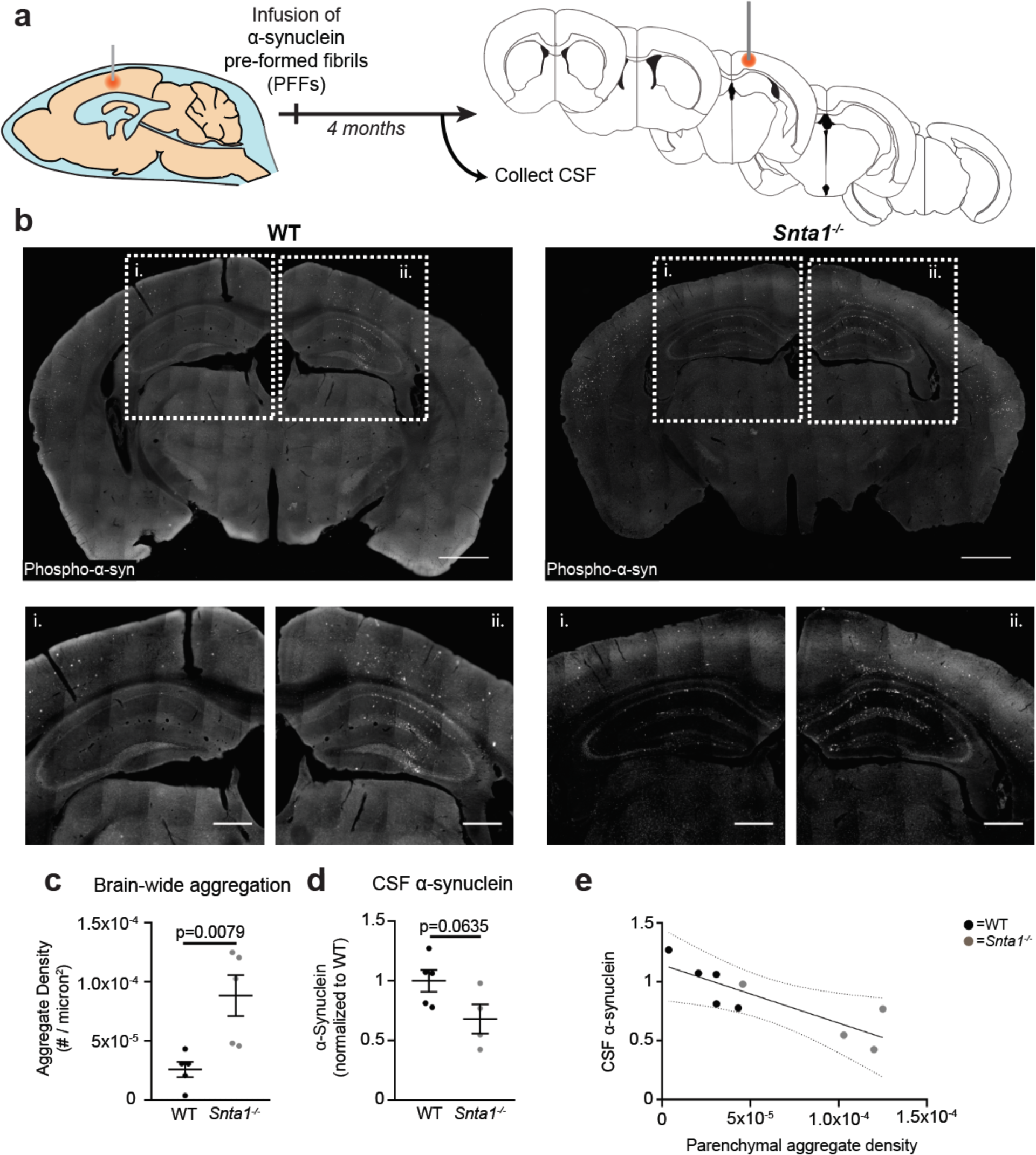
Insoluble α-synuclein aggregation is increased in *Snta1^-/-^* mice following inoculation with PFFs. (**a**) Schematic depiction of α-synuclein propagation studies following motor cortex injection of pre-formed fibrils (PFFs). The five coronal sections at which phospho-α-synuclein aggregate burden was evaluated 4 months following injection of PFFs at shown at right. Orange circle represents the site of infusion. (**b**) Representative images illustrating phospho-α-synuclein distribution in wild type and *Snta1^-/-^* mice. Scale bar = 1000 µm. Higher magnification images of contralateral (i) and ipsilateral (ii) cortex and hippocampus illustrate abundant aggregation in both regions. Scale bar = 200 µm. (**c**) Quantification of aggregate density determined by integration of phospho-α-synuclein aggregate counts across five sections reveals elevated global aggregate density in *Snta1^-/-^* mice (p=0.0079, Mann-Whitney U test, n=5/group). (**d**) A trend toward reduced CSF α-synuclein in *Snta1^-/-^* mice is demonstrated by ELISA-based quantification of CSF α-synuclein concentrations 4 months following PFF injection, normalized to mean wild type values (p=0.0635, Mann-Whitney U test. n=4-5/group). (**e**) Plotting aggregate density against CSF α-synuclein levels in individual animals reveals a linear negative association between these values across genotype. (p=0.005, R^2^=0.7063, Linear regression, n=4-5/group). For all plots, black bars represent the mean ± S.E.M.

## Discussion

While prior preclinical and postmortem studies have linked glymphatic dysfunction, changes in perivascular AQP4 localization, and the increased aggregation of AD-associated amyloid β and tau, the linkage between these features and α-synuclein aggregation, observed in PD and DLB, are less well defined. Emerging data suggest that in neurodegenerative conditions spanning AD, PD, DLB, Huntingtin’s disease, amyotrophic lateral sclerosis and Lewy body dementia, the protein aggregates undergo intercellular propagation ^59-63^. Yet whether host features, such as the impairment of glymphatic function with sleep disruption, aging, or injury, influence the rate of aggregate propagation remains unknown. Here, we demonstrate in a human postmortem case series that neocortical Lewy body pathology is associated with reduced perivascular AQP4 localization. In prior preclinical studies, we have demonstrated that loss of perivascular AQP4 localization impairs glymphatic function ^41,49^. In the present study, we report that soluble α-synuclein is cleared from brain tissue along perivascular pathways, while loss of perivascular AQP4 localization slows its clearance from the brain parenchyma to the CSF. Lastly, we demonstrate in a mouse model of α-synuclein preformed fibril injection that loss of perivascular AQP4 localization accelerates a-synuclein aggregate propagation. Based on these findings, we propose that loss of perivascular AQP4 localization contributes to the development and propagation of α-synuclein pathology in synucleinopathies including PD and DLB.

Previously, a human postmortem case series demonstrated a broad relationship between reduced perivascular AQP4 localization and amyloid β and tau aggregates in the setting of AD. In 2017, Zeppenfeld et al. reported that in individuals 33-105 years of age, perivascular AQP4 localization within the frontal cortical gray matter declined with age and was reduced in those with histopathologically-confirmed AD ^48^. Within this case series, reduced perivascular AQP4 localization was associated with both increased amyloid β plaque and tau pathological burden. A second, independent case series replicated these findings in individuals over 65 years of age, demonstrating that frontal cortical perivascular AQP4 localization was reduced in those with AD, while reduced perivascular AQP4 localization was associated with more severe cortical amyloid β plaque and tau pathological burden ^48^. Less is known concerning the relationship between α-synuclein pathology and changes in perivascular AQP4 localization in the human brain. A previous study examining human postmortem PD cases reported that α-synuclein pathology was associated with increased AQP4 expression in the temporal neocortex as a whole; however a negative correlation was observed between α-synuclein and AQP4 expression in neocortical layers V-VI ^64^. Notably, this study did not evaluate the perivascular localization of AQP4. In the present study, we demonstrate that in the human post-mortem brain neocortical α-synuclein pathology is associated with increased AQP4 expression at the pial surface and loss of perivascular AQP4 polarization. It is important to note that amyloid β, tau, α-synuclein and other pathological features are frequently comorbid within the aging population and that the relationships between these pathological features and AQP4 mis-localization may be complex. In the present study, we sought to avoid these confounds by selecting only cases with neocortical Lewy body pathology and excluding those with moderate to high CERAD neuritic plaque scores (2-3) or tau (Braak stages V-VI) pathology.

Although informative, these correlational histopathological studies cannot address whether the observed loss of perivascular AQP4 localization impairs glymphatic function, nor whether impairment of glymphatic function contributes to the development of amyloid β, tau or α-synuclein pathology in proteinopathies such as AD, PD or DLB. Similarly, these correlational histopathological studies also do not provide insight into whether sleep is a mediator/moderator of this relationship between AQP4 mis-localization and α-synuclein pathology. Prior studies in rodent models of aging, cerebrovascular injury, and traumatic brain injury demonstrated that these conditions each resulted in loss of perivascular AQP4 localization, which was in each case associated with impaired glymphatic function ^18,28,29,45-47,65^. Two subsequent studies utilizing *Snta1^-/-^*mice lacking perivascular AQP4 localization demonstrated that such mis-localization of AQP4 impairs glymphatic function ^39,49^. The present study extends these findings to the glymphatic clearance of α-synuclein. In the initial studies characterizing glymphatic function in rodents, we observed that both soluble amyloid β ^17^ and tau ^18^ are cleared from brain tissue along perivascular pathways. Herein we report that intraparenchymally-injected fluorescently-labeled α-synuclein is cleared through brain tissue along perivascular routes. In *Snta1^-/-^* mice, we observed that CSF levels of endogenous α-synuclein were lower than in wild type controls, while parenchymal levels of α-synuclein remained elevated. These observation suggest that the glymphatic clearance of α-synuclein from brain tissue into the CSF compartment is impaired with reduced perivascular AQP4 localization.

Prior studies that have implicated AQP4 in glymphatic pathway function have generally relied upon global *Aqp4* knockout lines ^17,39^, as have studies demonstrating that impairment of glymphatic function exacerbates protein mis-aggregation in the brain. Deletion of the *Aqp4* gene in mouse models has been shown to exacerbate amyloid β pathology ^40,41^ and tau pathology ^18,43^. Focusing on synucleinopathy, a recent study reported impaired clearance of injected α-synuclein in the setting of *Aqp4* gene deletion. Genetic deletion of *Aqp4* exacerbated α-synuclein pathology in mice overexpressing human A53T-α-syn, and overexpression of human A53T-α-syn decreased polarization of AQP4 and impaired glymphatic function ^66^. A similar study reported increased deposition of injected α-synuclein PFFs and decreased glymphatic clearance in AQP4^+/-^ heterozygote mice ^67^. While these studies suggest a key role of AQP4 in regulating α-synuclein aggregation and aggregate propagation, they are limited by the fact that they do not recapitulate the AQP4 phenotype, including reduced perivascular AQP4 localization, observed in the aging and injured brain. In our prior studies we have observed that loss of perivascular AQP4 localization in the *Snta1^-/-^* mouse exacerbates amyloid β pathology ^41,49^. In the present study we extend these observations to the setting of synucleinopathy, observing that *Snta1* gene deletion exacerbates α-synuclein aggregate propagation.

The present study is distinct from most prior studies of the effect of glymphatic disruption on protein mis-aggregation in utilizing an α-synuclein PFF aggregate propagation assay on a wild type background rather than evaluating what is likely to be cell-autonomous aggregate development in genetically-driven models. In the APP/PS1^40^, Tg2576^49^, and 5XFAD^41^ amyloidosis lines, the PS19 tauopathy line^43^, and the A53T-α-syn synucleinopathy lines^66^, mutant proteins are expressed widely and pathological burden may develop in individual cells solely by virtue of their intrinsic expression of the pro-fibrillary forms of these proteins. In contrast, in the present model α-synuclein PFFs are injected locally into a wild type α-synuclein background, and the spread of pathologically phosphorylated α-synuclein to other remote regions can only occur through the cell-to-cell propagation of pathological fibrils, and through the recruitment of non-pathological wild type mouse α-synuclein into pathological α-synuclein aggregates detected by P-Syn antibodies. The present study suggests that impairment of glymphatic function through loss of perivascular AQP4 localization does not simply promote general protein mis-aggregation, but also exacerbates the process of cell-to-cell aggregate propagation. Whether these effects extend beyond the propagation of α-synuclein aggregates, to the propagation of other pathological proteins, such as pathological tau aggregates, remains an important question to address in the future.

The glymphatic clearance of interstitial solutes is more rapid during sleep ^5,21^, and is driven by synchronous electrical activity characteristic of different sleep stages, such as the delta band activity observed during slow wave sleep ^68,69^. Aging, neurodegenerative conditions, cerebrovascular disease, and traumatic brain injury are each clinically associated with significant sleep disruption often occurring early on in the pathogenesis of the disease course ^1,70,71^, in addition to causing glymphatic dysfunction in preclinical models ^18,28,45-47^. Thus, glymphatic function may be impaired directly, including through loss of perivascular AQP4 localization, as well as indirectly, as sleep disruption reduces the amount of time spent under conditions that promote more rapid glymphatic clearance. There may also be a bidirectional relationship between mis-localization of AQP4 and sleep disruption; indeed, recent studies have found that sleep deprivation and disruption in mice caused mis-localization of AQP4 at the astrocytic endfeet ^65,72^. Notably, another study reported changes in AQP4 expression following sleep fragmentation in wild-type and 5XFAD mice, though AQP4 localization was not investigated ^73^. It has not been reported whether AQP4^-/-^ or Snta1^-/-^ mice demonstrate changes in sleep, such studies may provide evidence for the possible affects of AQP4 expression and localization on sleep. Furthermore, sleep disruption exhibits a specific association with synucleinopathies that should be also be considered ^6,34,35,74,75^. Indeed, a recent meta-analysis found that >96% of individuals with RBD phenoconverted to overt synucleinopathy (principally PD or DLB) within 14 years of diagnosis of this violent and disruptive sleep disorder ^74^. Thus, relationships between AQP4 mis-localization and sleep disruption in contributing to glymphatic impairment in synucleinopathies, and other neurodegenerative conditions, are likely to be highly intertwined.

This study had a number of limitations that we would like to acknowledge. First, given the cross-sectional nature of the human post-mortem studies, it was not possible to assess the directionality of the association between AQP4 mis-localization and Lewy body pathology. However, our results using experimental manipulations in mice suggest that AQP4 mis-localization is capable of promoting α-synuclein pathology under experimental conditions. However, further studies should confirm this relationship in the human brain. These studies did not formally assess sleep, although it is well established that glymphatic biology is more active during sleep and that sleep disruption is commonly observed in both human and mouse synucleionpathy. Sleep may be a mediator/moderator of the relationship between AQP4 mis-localization and α-synuclein pathology, as well as directly affect AQP4 localization and α-synuclein pathology via other pathways outside the glymphatic system. Future research is needed to fully flesh out the relationships between sleep, AQP4, and synuclein biology in mice and humans.

In conclusion, we demonstrate that human neocortical Lewy body pathology is associated with a loss of perivascular AQP4 localization. Such mis-localization of AQP4 in animal models impairs glymphatic function, including the clearance of α-synuclein from brain tissue to the CSF. We also show that these changes are sufficient to promote the propagation of pathological α-synuclein aggegates. In total, these findings suggest that glymphatic impairment may contribute to the development and progression of synucleinopathies such as PD and DLB, and may provide insight into the clinical linkage between sleep disruption and synucleinopathy.

## Acknowledgements

We thank Mick Metcalf for his contributions to ROI generation and the OHSU Advance Light Microscopy (ALM) Core for their assistance with image acquisition troubleshooting. We also thank Ms. Aimnee Schantz and Mr. John Campos for outstanding administrative support, and the brain donors and their families. This study was supported by grants from the National Institute of Health awarded to J.J. Iliff (R01-NS089709 and R01-AG054456), M.J. Simon (F31-AG054093) and Dr. Sue Aicher and the ALM core (P30-NS061800); and Veterans Affairs funding to J.E. Elliott (RX002947). Post-mortem human samples used in this study were obtained from the University of Washington UW Biorepository and Integrated Neuropathology (BRaIN) Laboratory, which is supported by the Alzheimer’s Disease Research Center (AG066509), the Adult Changes in Thought Study (AG006781),

