## Supplementary Figures & Tables for "Aquaporin-4 mis-localization slows glymphatic clearance of α-synuclein and promotes α-synuclein pathology and aggregate propagation"

### Supplementary Tables & Figures

#### Supplementary Table 1: Individual demographic and neuropathological data for human post-mortem tissue

| Lewy Body Status | Sample ID | Age (yrs) | Sex | LBD Diagnosis | Braak Stage | CERAD Score |
|---|---|---|---|---|---|---|
| Control | 1 | 77 | M | None | II | Sparse (1) |
| Control | 2 | 80 | M | None | I | Absent (0) |
| Control | 3 | 82 | F | None | III | Sparse (1) |
| Control | 4 | 88 | F | None | IV | Absent (0) |
| Control | 5 | 91 | F | None | II | Sparse (1) |
| Control | 6 | 100 | F | None | IV | Sparse (1) |
| Control | 7 | 77 | F | None | II | Sparse (1) |
| Control | 8 | 78 | M | None | I | Absent (0) |
| Control | 9 | 79 | M | None | II | Absent (0) |
| Control | 10 | 86 | M | None | III | Sparse (1) |
| Control | 11 | 86 | M | None | II | Sparse (1) |
| Control | 12 | 87 | M | None | IV | Sparse (1) |
| Control | 13 | 87 | M | None | II | Sparse (1) |
| Control | 14 | 88 | F | None | IV | Absent (0) |
| Control | 15 | 89 | F | None | II | Sparse (1) |
| Control | 16 | 89 | F | None | II | Sparse (1) |
| Control | 17 | 91 | F | None | IV | Absent (0) |
| Control | 18 | 84 | M | None | II | Absent (0) |
| Control | 19 | 82 | M | None | IV | Sparse (1) |
| Case | 1 | 76 | F | Neocortical | II | Sparse (1) |
| Case | 2 | 88 | F | Neocortical | I | Sparse (1) |
| Case | 3 | 70 | M | Neocortical | II | Sparse (1) |
| Case | 4 | 91 | F | Neocortical | II | Sparse (1) |
| Case | 5 | 73 | F | Neocortical | II | Absent (0) |
| Case | 6 | 70 | M | Neocortical | I | Sparse (1) |
| Case | 7 | 80 | M | Neocortical | IV | Sparse (1) |
| Case | 8 | 69 | M | Neocortical | III | Sparse (1) |
| Case | 9 | 87 | M | Neocortical | I | Sparse (1) |
| Case | 10 | 82 | F | Neocortical | III | Sparse (1) |
| Case | 11 | 96 | F | Neocortical | IV | Absent (0) |
| Case | 12 | 62 | M | Neocortical | III | Sparse (1) |
| Case | 13 | 75 | M | Neocortical | III | Absent (0) |
| Case | 14 | 91 | F | Neocortical | I | Absent (0) |
| Case | 15 | 84 | M | Neocortical | IV | Sparse (1) |
| Case | 16 | 98 | F | Neocortical | III | Absent (0) |
| Case | 17 | 77 | M | Neocortical | IV | Sparse (1) |
| Case | 18 | 88 | M | Neocortical | IV | Sparse (1) |
| Case | 19 | 96 | F | Neocortical | IV | Sparse (1) |
| Case | 20 | 85 | M | Neocortical | IV | Sparse (1) |
| Case | 21 | 93 | F | Neocortical | IV | Sparse (1) |

**Supplemental Figure 1**

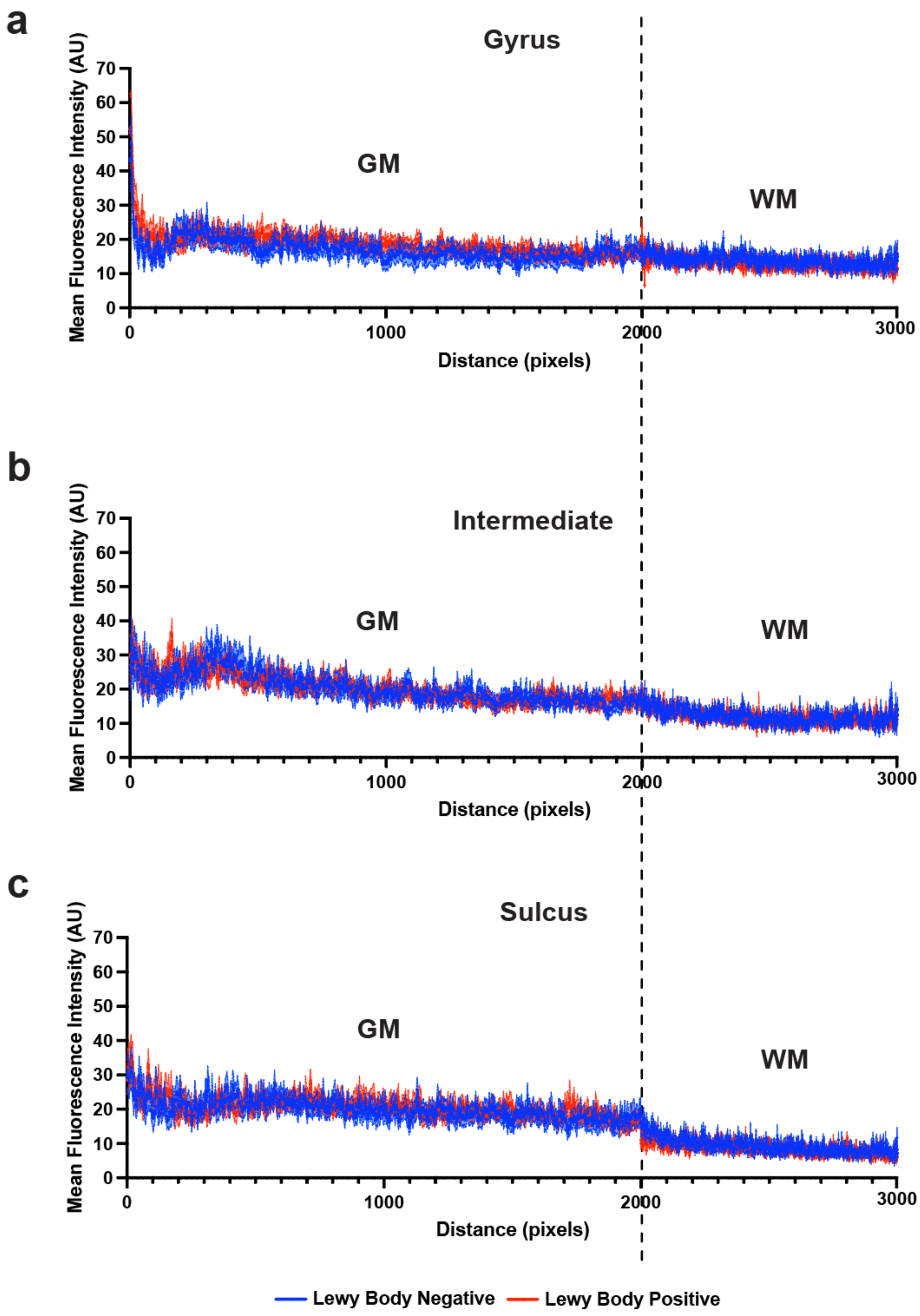

**Supplemental Figure 1: Intensity projection plots of AQP4 fluorescence across human post-mortem brain tissue with and without α-synuclein pathology.** Three ROI lines were drawn at the (**a**) gyral crest, (**b**) midway down the gyrus, and at the (**c**) sulcal depth to measure AQP4 fluorescence across the tissue. Fluorescence intensity plots were averaged for each region and then averaged across groups. Intensity plots extended from the pial surface and through the grey matter and adjoining lines extended through the white matter. For statistical analysis, gray matter (GM) line segments were divided into 3 equal segments corresponding to the superficial, middle, and deep GM regions of the laminar shell analysis. White matter (WM) line segments were divided into 2 equal segments corresponding to superficial and deep WM. (**a**) At the crest of the gyrus, main effects of depth ($p<0.0001$) and depth x group interactions ($p=0.0197$) in the superficial GM, main effects of depth ($p<0.0001$) in the middle GM and superficial WM, and a main effect of depth x group interaction ($p<0.0001$) in the deep GM were detected. (**b**) Midway between the crest of the gyrus and depth of the sulcus, main effects of depth ($p=0.0485$) and depth x group interactions ($p=0.0227$) in the superficial GM, main effects of depth ($p<0.0001$) in the middle GM and superficial WM, and a main effect of depth x group interaction in the deep GM ($p=0.0232$) and deep WM ($p<0.0001$) were detected. (**c**) At the sulcus, main effects of depth ($p<0.0001$) in the superficial GM and middle GM, main effects of depth ($p<0.0001$) and depth x group interaction ($p=0.0010$) in the deep GM, and main effect of depth in the superficial WM ($p<0.0001$) and deep WM ($p=0.0110$) were detected**.** Data were analyzed by 2-way repeated measures ANOVA followed by Sidak's post-hoc test and are mean ± SEM from $n$ = 19-21 per group.

**Supplemental Figure 2**

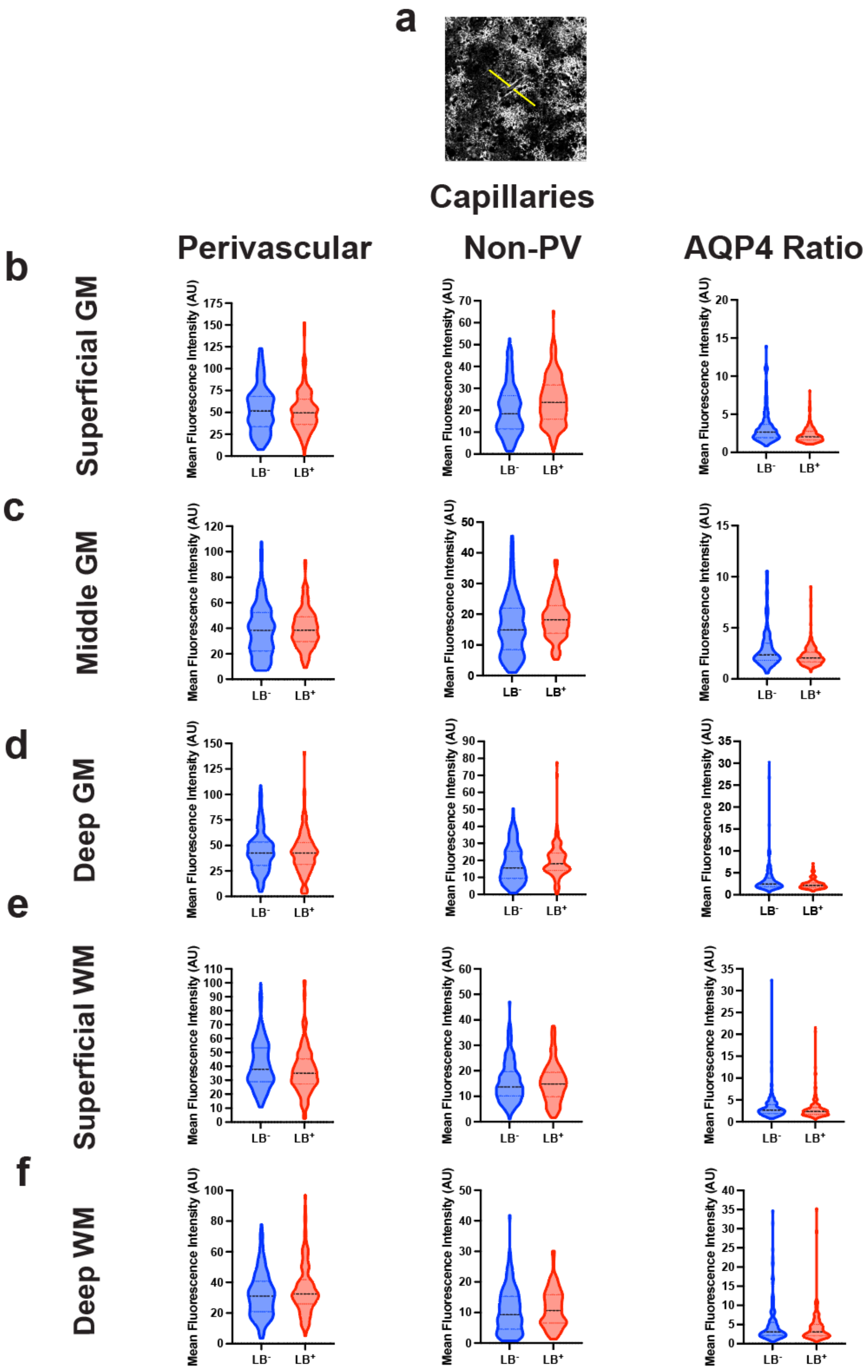

**Supplemental Figure 2: Data distribution of perivascular, non-perivascular, and AQP4 polarization ratio of capillaries within cortical laminae.** (**a**) Representative image of capillary with drawn yellow lines indicating intensity projection plots drawn through the perivascular endfoot, continuing through the surrounding astrocytes and neuropil. The pixel intensities were binned and averaged across these line segments. The first 3 pixels were averaged for the perivascular endfoot segment, the last 20 pixels were averaged for the non-perivascular segment. AQP4 surrounding capillaries were quantified in (**b**) superficial grey matter (GM) (**c**) middle GM (**d**) deep GM (**e**) superficial WM and (**f**) deep WM. In the violin plots, the central line represents the median, upper line represents the upper interquartile range (IQR), and the lower line represents the lower IQR. AQP4 was measured around an average of 10 capillaries per region per subject from *n* = 19-21 subjects per group.

**Supplemental Figure 3**

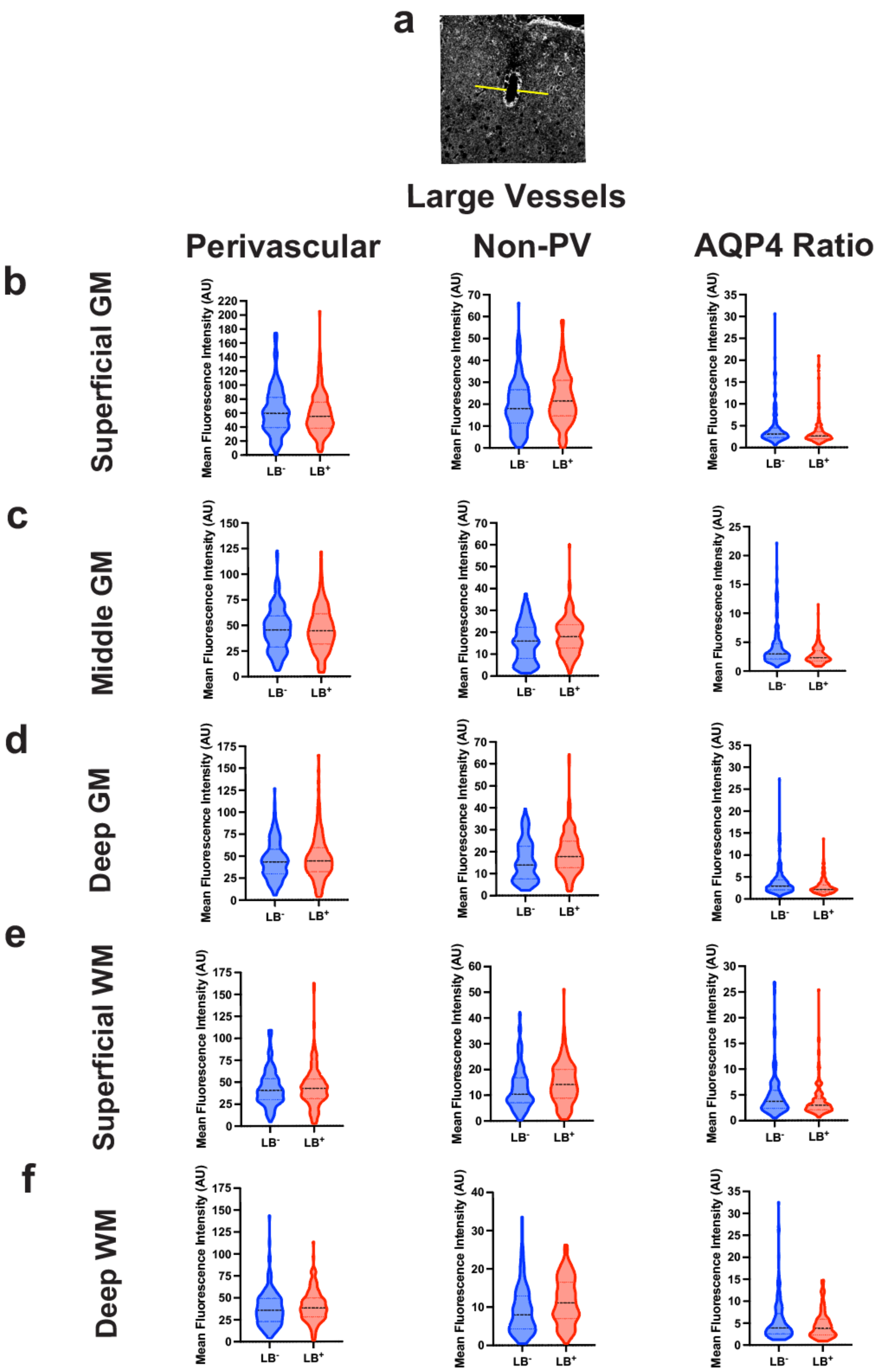

**Supplemental Figure 3: Data distribution of perivascular, non-perivascular, and AQP4 polarization ratio of large vessels within cortical laminae.** (**a**) Representative image of large vessel with drawn yellow lines indicating intensity projection plots drawn through the perivascular endfoot, continuing through the surrounding astrocytes and neuropil. The pixel intensities were binned and averaged across these line segments. The first 3 pixels were averaged for the perivascular endfoot segment, the last 20 pixels were averaged for the non-perivascular segment. AQP4 surrounding large vessels were quantified in (**b**) superficial grey matter (GM), (**c**) middle GM, (**d**) deep GM, (**e**) superficial WM, and (**f**) deep WM. In the violin plots, the central line represents the median, upper line represents the upper interquartile range (IQR), and the lower line represents the lower IQR. AQP4 was measured around an average of 10 large vessels per region per subject from *n* = 19-21 subjects per group.

**Supplemental Figure 4**

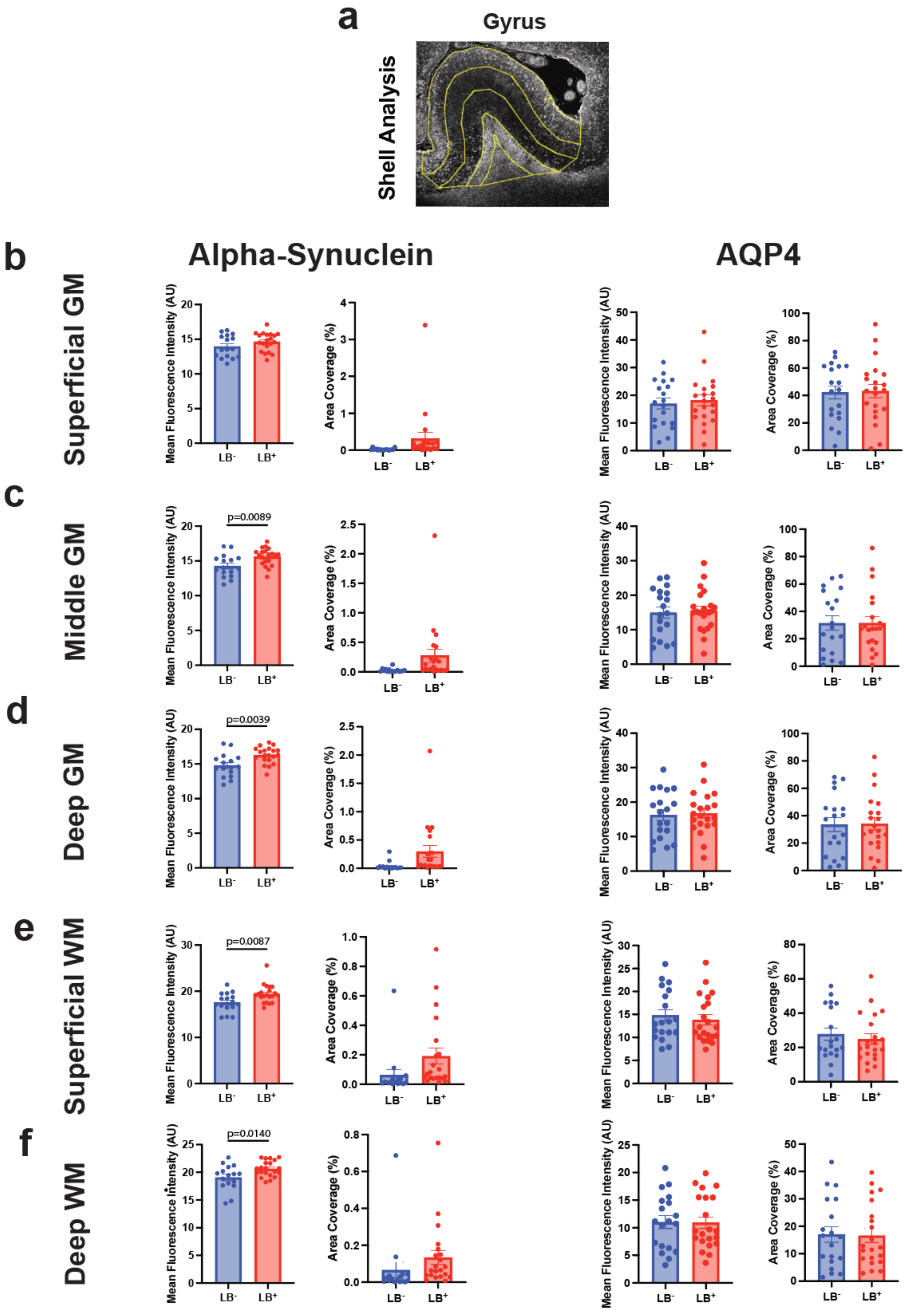

**Supplemental Figure 4: Increased α-synuclein pathology and unchanged AQP4 expression as assessed by laminar shell analysis.** (**a**) Representative image of laminar shell regions of interest (ROI) starting with the outer (**b**) Superficial Gray Matter (GM), (**c**) Middle GM, (**d**) Deep GM, (**e**) Superficial WM, and the innermost (**f**) Deep WM ROI. Quantification of phosphorylated α-synuclein (p-S129) and AQP4 immunoreactivity within these ROIs is shown, including fluorescence intensity (left columns) and area coverage (right columns). Phosphorylated α-synuclein levels were significantly increased in the Middle GM ($p=0.0089$), Deep GM ($p=0.0039$), Superficial WM ($p=0.0087$), and Deep WM ($p=0.0140$) in Lewy Body Positive cases compared to controls while no differences were seen in overall AQP4. Data are mean ± SEM from n= 19-21 subjects/group and were analyzed by unpaired t-test.

**Supplemental Figure 5**

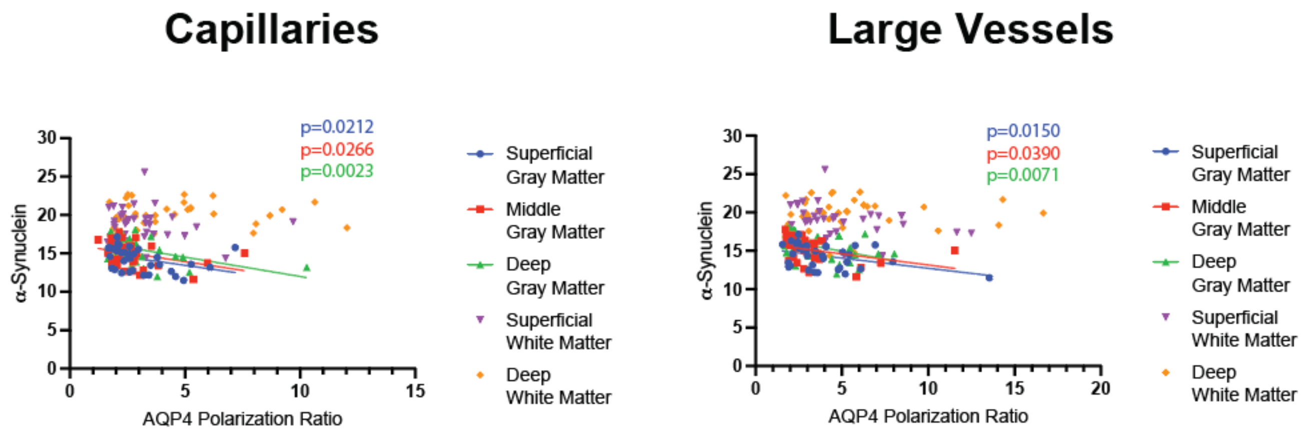


**Supplemental Figure 5: Association of α-synuclein pathology and lewy body status with AQP4 polarization ratio.** Linear regression analyses were performed to assess the relationship between α-synuclein and AQP4 localization surrounding capillaries and large vessels. Averaged values of α-synuclein fluorescence and AQP4 polarization ratio within laminar regions were assessed. A significant negative relationship between α-synuclein and AQP4 polarization ratio was observed surrounding capillaries (caps) and large vessels (LVs) in the Superficial Gray Matter (GM) (caps: p=0.0212, $R^2$=0.1465; LVs: p=0.0150, $R^2$=0.1618), Middle GM (caps: p=0.0266, $R^2$=0.1403; LVs: p=0.0390, $R^2$=0.1303), and Deep GM (caps: p=0.0023, $R^2$=0.2419; LVs p=0.0071, $R^2$=0.1943).

**Supplemental Figure 6**

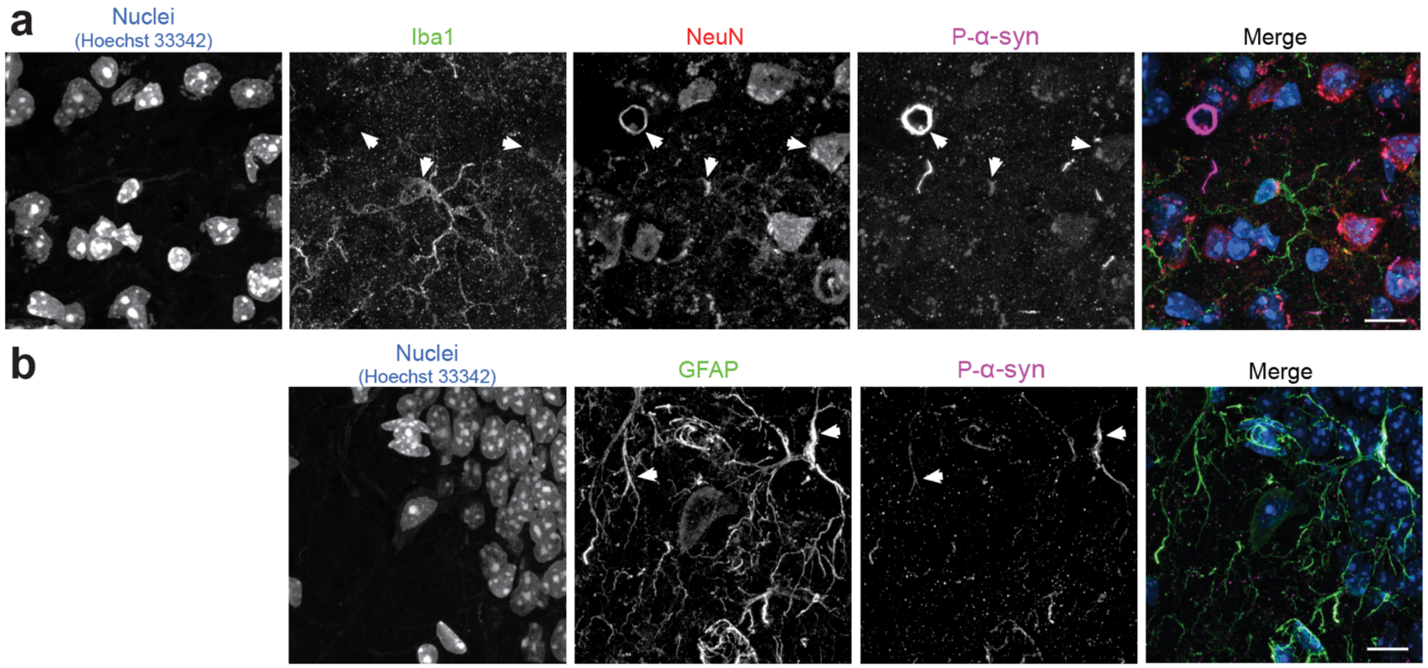


**Supplemental Figure 6: Cell types exhibiting α-synuclein aggregate formation following PFF injection**. (**A**) Immunofluorescence imaging of phospho-α-synuclein (magenta) with cell type markers for microglia (Iba1, green) and neurons (NeuN, red) reveal colocalization with both cell types, though more abundantly with neurons. (**B**) Immunofluorescence imaging of phospho-α-synuclein an astrocytic marker (GFAP, green) reveals colocalization. Arrows indicate sites of co-localization as well as phospho-α-synuclein exhibiting various morphologies (neuritic, glial and somatic inclusions as well as diffuse). Merged images at left. Scale bar = 10μm.
